# GlyComboCLI enables command line-based FAIR workflows for glycan composition assignment in mass spectrometry data

**DOI:** 10.64898/2026.05.13.725058

**Authors:** Maia I. Kelly, W. C. Mike Thang, C. N. Ignatius Pang, O. Johan R. Gustafsson, Christopher Ashwood

## Abstract

Glycans are integral biomolecules whose presence cannot be predicted from genomic data alone, necessitating experimental characterisation through approaches including mass spectrometry. Assignment of glycan compositions to observed mass to charge ratios is computationally challenging due to the potential monosaccharide diversity and existing tools lack the required flexibility for integration into automated bioinformatic workflows.

Here, we present GlyComboCLI, an open-source command-line application for the assignment of glycan compositions to mass spectrometry data which expands upon the previous GUI application, GlyCombo. GlyComboCLI accepts mass lists and vendor-neutral mzML files, supports diverse monosaccharides, derivatisation states, reducing-end modifications and adducts, and is validated through automated tests spanning supported search parameters, input formats, and instrument vendors. Outputs are compatible with downstream tools including Skyline and GlycoWorkBench while GlyTouCan accessions support persistent identification of registered base compositions. Deployment as a standalone executable, a Docker container, and a Galaxy tool, supports FAIR and reproducible workflows.

Applied to published mouse and human glycomics datasets, GlyComboCLI reproduced major qualitative glycomic motifs and relative abundance trends, while Galaxy workflow reproduced expected effects of sialidase treatment. These results demonstrate a flexible, scalable, and reproducible approach for glycan composition assignment within automated glycomics workflows.

## Introduction

Glycobiology is the study of products of protein glycosylation called glycans, which includes oligosaccharides and polysaccharides, and these carbohydrates play key roles in the folding and structural stability of proteins^1^, protein trafficking^2^, cell adhesion^3^ and cellular communication^4^. Unlike proteins or nucleic acids, glycans are not template-derived, meaning that they cannot be predicted solely from the genome. They also reflect an organism’s environment, including glycome remodelling in response to infection or cell status^5,6^. For these reasons the overall landscape of these glycans, the glycome, must be defined experimentally.

A variety of analytical approaches can be employed to characterise and quantify the glycans in a given sample, including fluorophore-based techniques^7,8^ and mass spectrometry (MS)^9,10^. Unlike fluorophore-based techniques, MS directly measures intact glycans through their mass to charge ratio (m/z), which reflects the composite sugars (i.e. composition). Subsequently, specific glycans of interest can be subjected to fragmentation and detection (known as MS2 or MS/MS) to verify the predicted monosaccharide composition and clarify how the composite sugars are arranged (i.e. structure). One unique approach, OxoPlot^11^, prioritises structural epitope elucidation by assigning structural features to specific m/z values observed in MS2 spectra, subsequently followed by composition assignment.

Bioinformatic workflows can increase high-throughput analysis by increasing automation, reproducibility, and in turn decreasing manual, time consuming interpretation. The software-based assignment of glycan compositions to observed m/z values is often the first step in the largely manual structural analysis workflow. This is necessary because, unlike proteins, glycans can exhibit a high degree of structural and compositional diversity due to the numerous types of monosaccharides, branching patterns, and additional chemical modifications^12^. As the number of combinations of chained and branched monosaccharides, chemical modifications, and glycan sizes increases, the assignment of glycans to their observed mass-to-charge ratios becomes computationally challenging, highlighting the potential benefits of tailored automated workflows.

A seminal software tool for glycan composition assignment is GlycoMod^13^. The software enables users to match observed glycan m/z values with potential glycan compositions from a pre-selected monosaccharide search range. Unfortunately, the software is closed source, which can restrict the integration of GlycoMod with bioinformatic workflows. In addition, only text/integer input is supported, preventing continuous workflows. GlyCombo^14^ was developed to support the scalable, integrative assignment of glycan composition to masses (either as text or mzML^15^ input) within bioinformatic workflows and has been utilised to characterise the glycome in colorectal cancer^16^, mucin^17^, and serum^18^ contexts. In its initial version, the graphical user interface (GUI) format made analysis user-friendly and accessible. However, this is a bottleneck for automated workflows and web server deployments, both of which require command-line operation.

GlyComboCLI returns all compositions within a user-defined search space, whose calculated precursor masses satisfy the specific mass-error tolerance. Computational validation therefore tests whether the expected mass-matched compositions are returned across supported search parameters and input formats. Experimental ambiguities, including incorrect monoisotopic precursor assignment, or multiple compositional candidates for a given precursor mass (isomers), require downstream evaluation using precursor isotopic distribution, MS2 annotation, or other orthogonal evidence. Accordingly, GlyComboCLI was evaluated using both automated software tests and published glycomics datasets.

In this work, we present a substantive update with GlyComboCLI, a command-line program for assigning glycan compositions to mass spectrometry data. GlyComboCLI is open source and open access, and aligns with the findable, accessible, interoperable, and reusable (FAIR) principles^19^. It is available as both a command-line and a container-based application. This ensures that the software can be reused in real world analytical scenarios, including as a local work station computation, where it is a middle step in a pipeline, and as a Galaxy tool^20^, which can be used in a modular fashion for reproducible data analysis workflows on shared compute systems. The ready availability and flexibility of GlyComboCLI allows the software to fit into custom workflows required by research scientists, regardless of where they need to undertake these analyses.

## Experimental section

### GlyComboCLI development

GlyComboCLI (1.0.0) is a command line interface application developed as a complement to the GlyCombo application available as a graphical user interface. GlyComboCLI was developed in C# using .NET 8 from a single cross-platform codebase and distributed as a portable .NET 8 package for Windows/macOS/Linux plus a Windows x64 executable package. The open-source code, under the Apache 2.0 license, is publicly available on GitHub (https://github.com/Protea-Glycosciences/GlyComboCLI) and archived on Zenodo (https://doi.org/10.5281/zenodo.20063128), with software metadata registered at bio.tools^21^ (https://bio.tools/glycombocli). Versioned GlyComboCLI container images are available from Docker Hub at https://hub.docker.com/r/proteaglycosciences/glycombo/ The Galaxy wrapper source is available at https://github.com/Protea-Glycosciences/GlyComboGalaxy and the validated Galaxy tool is distributed through the Main Galaxy ToolShed at https://toolshed.g2.bx.psu.edu/repos/proteaglycosciences/glycombo.

GlyComboCLI accepts two input formats, .txt and .mzML, specified with the --file flag. The text format allows for a plain-text mass list of user defined precursor masses for compositional matching. The mzML file is vendor neutral with GlyComboCLI directly extracting the precursor m/z values, charge state and polarity. Glycan compositions are prepared from combinatorial search from each user defined precursor and search range parameters. Monosaccharide support is available for hexose, *N*-acetylhexosamine, deoxyhexose, hexuronic acid, hexosamine, pentose, 2-keto-3-deoxy-D-glycero-D-galacto-nononic acid, *N*-acetylneuraminic acid (NeuAc), *N*-glycolylneuraminic acid (NeuGc), and *N-* acetyldeoxyhexose. Additional modified sialic acid species, lactonised, ethyl esterified, dimethylamidated, and ammonia amidated forms of NeuAc and NeuGc are available along with acetylation, phosphorylation and sulfation. Five custom monosaccharides are available for user definition through elemental composition (carbon, hydrogen, nitrogen, and oxygen) with a mass value.

Through flags in GlyComboCLI, users are able to define derivatization state (native, permethylated or peracetylated), reducing end modifications (free, reduced, InstantPC, Rapifluor-MS, 2-aminobenzoic acid, 2-aminobenzamide, procainamide, Girard’s reagent P, and custom reducing ends) and adducts (neutral, [M+H]+, [M+Na]+, [M+NH4]+, [M+H]-, [M+FA]-, [M+AA]-, [M+TFA]-, and custom adducts). Further user options include mass error, defined in either Daltons (Da) or parts per million (ppm) and optional off-by-one search.

GlyComboCLI performs glycan composition assignment based on precursor *m/z*, the user-defined search space, and mass-error tolerance, but does not perform internal quality control filtering. In the Galaxy workflow, filtering and QC visualisation were performed by independent downstream workflow nodes using the GlyComboCLI output, functions not implemented within GlyComboCLI. An output schematic is presented in Figure S1. For previously reported glycan compositions in GlyTouCan, accessions are provided as a separate column in the output by matching between the GlyComboCLI composition output and a statically bundled reference of the GlyCosmos glycans list (https://download.glycosmos.org/glycosmos_glycans_list.csv, latest snapshot 2026-11-02).

Upon completion of the search, GlyComboCLI produces up to three output files, depending on the input file. A text input file will yield two output files, a parameters text file with all user defined options and a csv results file containing all possible glycan compositions within the search space. An mzML file will yield three output files, the two previously mentioned files, with an additional csv file for direct import into Skyline. Parameter files are available for cross-platform use with GlyCombo GUI.

### Local GlyComboCLI application

After downloading the raw files corresponding to GPST000289, GPST000398, and GPST000338, msconvert (v3.0.25316) converted the raw vendor files to .mzML, only retaining MS2 scans, with 4 files converted in parallel in a single instance. Through the use of a parallelised batch file, four concurrent GlyComboCLI instances assigned glycan compositions to the observed masses (Settings: hMin 2, hMax 12, nMin 2, nMax 8, fMax 3, sMax 5, gMax 5, Derivatisation: Native, Reducing end: Reduced, Mass error: 15 ppm). For the human serum N-glycan study, the GlyComboCLI settings were as follows: hMin 2, hMax 9, nMin 2, nMax 6, fMax 3, sMax 4, Derivatisation: Native, Reducing end: Reduced, Mass error: 10 ppm.

Following GlyComboCLI composition assignment, quality control was performed downstream. In Skyline-daily (v26.1)[22], [23], [24], precursor isotope distributions were assessed using three extracted isotopic peaks from the supplied molecular formula from GlyComboCLI, and candidates with an isotopic dot product (idotp) below 0.9 were excluded to remove poor-quality MS1 matches and monoisotopic precursor assignment errors. Composition-based filtering was performed by a PowerShell script to only include compositions that met the following rule (NeuAc/NeuGc ≤ Hex − 3). These filtering steps are external to GlyComboCLI. GlycoWorkBench[25] annotated MS2 which were assigned glycan compositions by GlyCombo. B/Y/C/Z fragment types[26] were enabled for annotation with a maximum of three cleavages, using a 25 ppm mass accuracy threshold.

### Docker Image Construction of GlyCombo

To improve cross-platform availability and reproducibility across environments, GlyComboCLI was additionally made into a Docker container and publicly hosted on Docker Hub. Docker packages the application and environment into a self-contained image available for users to quickly run on Windows, macOS, and Linux systems. A version-controlled Dockerfile is maintained within the GlyComboCLI source repository. For each GlyComboCLI release, GitHub Actions builds the Docker image from the corresponding source code and validates it through automated command-line and Galaxy integration tests before publication to Docker Hub. All command line flags are identical to those used in the previously described CLI executable. Input and output files utilize a mounted directory.

### GlyCombo Integration in Galaxy

The GlyCombo Docker image was utilized for integration with Galaxy^22^, contribution to the Galaxy ToolShed^23^, and deployment to the Galaxy Australia service to enable access and easy integration into existing Galaxy workflows. Galaxy Australia is an open and accessible cloud platform that is built on a national distributed network of underpinning compute, and which is based on the global Galaxy platform and project^24^. Galaxy Australia therefore provides an ideal platform to make GlyComboCLI available to a broad life sciences audience, and in particular those researchers who do not have the time or incentive to upskill in bioinformatics.

All previously mentioned user defined options were maintained for the integration with Galaxy, which replaces command line flags with a web-based GUI. To maintain reproducibility and robustness of the software, and to ensure all features are preserved, GlyComboCLI is covered by 17 automated xUnit tests, while the Galaxy XML wrapper contains 21 embedded functional tests. The Galaxy tests exercise text and mzML inputs, instrument-vendor-derived mzML files, search parameters, and custom monosaccharide, reducing-end and adduct functionality. The wrapper is tested against the GlyComboCLI Docker container using Planemo before release. All vendor files were converted into .mzML format using msconvert (v3.0.25316)^25,26^.

### GlyCombo Galaxy application

GlyCombo Galaxy was evaluated in the context of a comparison between control and sialidase treated N-glycans from Panorama Public (https://panoramaweb.org/GlycoMS2Diagnostics.url)^27^. The Galaxy workflow (https://doi.org/10.48546/WORKFLOWHUB.WORKFLOW.2167.1) included raw file conversion (via msconvert), GlyComboCLI, filtering of the GlyCombo output, and visualisation of results through QC plots and summary plots (scatterplots and heatmaps via ggplot2 and ggplot, respectively) which were subsequently imported into figure panels.

## Results

### An open-source command-line version facilitates a breadth of informatic approaches

GlyCombo, a Windows-based GUI application was revised into a command line interface (CLI) to generate GlyComboCLI, enabling local use (e.g. Windows laptop) and online use through web instances and cloud-based platforms (*e*.*g*. Galaxy) (**Fig 1A**). In its local form, GlyComboCLI can serve as an intermediate step within a single piped command that spans multiple operations. One such example is shown in **Fig 1B**, where msconvert (mzML conversion), GlyComboCLI (glycan composition assignment), and Skyline (glycan verification) are run in series to output a list of glycan compositions identified in a given raw file. This list can then be subsequently used in orthogonal software such as GlycoWorkBench^28^ for MS2 verification of compositions.

**Figure 1.**
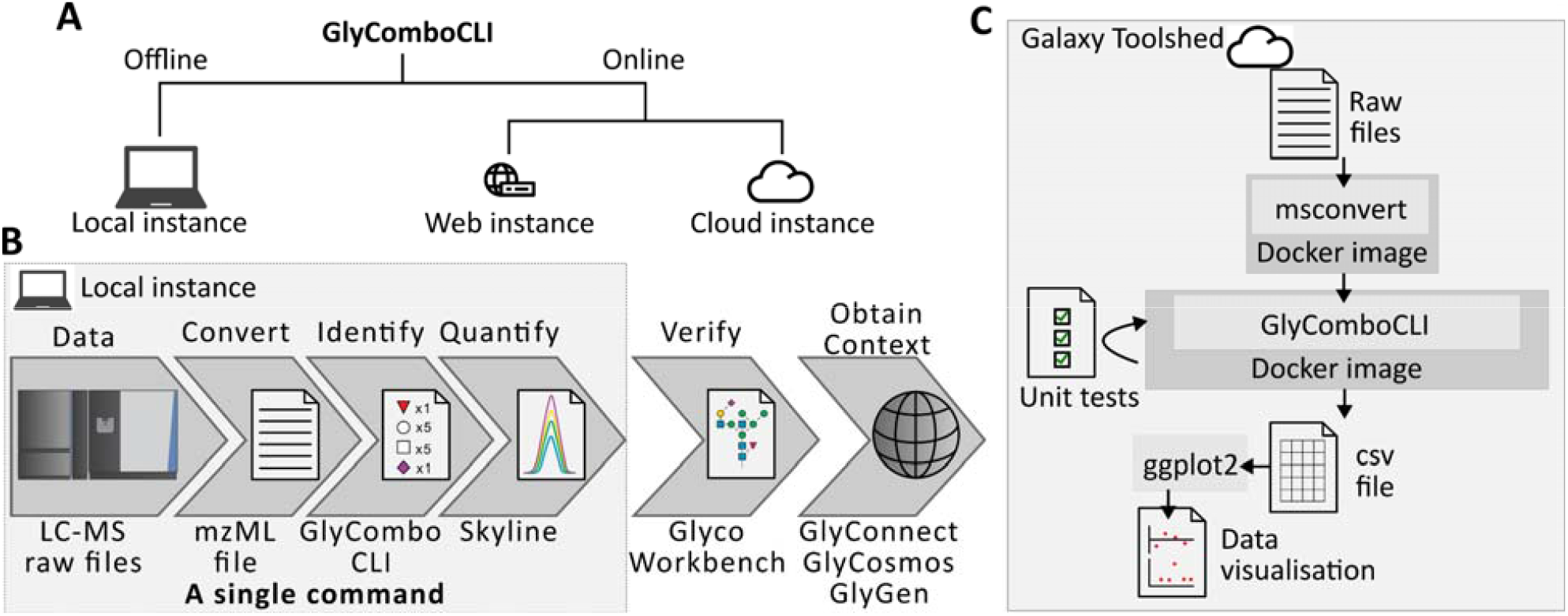
An open-source command line version of GlyCombo enables FAIR workflows for glycomics. **A** Three flavours of GlyCombo availability are enabled by GlyComboCLI. **B** A semi-automated workflow for data analysis of glycomics data. **C** Online versions of GlyComboCLI including Galaxy ToolShed are enabled by a Docker image.

When an exact registered base composition is available in the locally-stored GlyCosmos glycan database, GlyComboCLI additionally reports the corresponding GlyTouCan accession^29,30^, providing a persistent identifier for downstream reuse. Following composition assignment, results can be further contextualised using community glycoinformatics resources: GlyCosmos^31^ integrates glycan information across a broader network of glycoscience resources, GlyGen^32^ provides organism and biosynthetic context (related compositions), and GlyConnect Compozitor ^33,34^ enables relationships among sets of observed glycan compositions to be explored. Thus the “obtain context” step in **Fig 1B** represents post-assignment biological interpretation rather than an additional GlyComboCLI processing step.

Online web servers and cloud platforms improve accessibility and reproducibility by enabling researchers to run analysis on shared infrastructure through a browser. Using a Debian Linux image containing GlyComboCLI, the same glycan assignment process can be performed reproducibly through Galaxy (**Fig 1C**). Within Galaxy, additional tools, including msconvert and ggplot2, can be incorporated into the same workflow to further enhance reproducibility while embedded integration tests verify consistency across GlyComboCLI versions.

### Local GlyComboCLI use for high-throughput glycan assignment and quantification

A prior study analysed N-glycans released from 26 tissue locations across two mice using diagnostic fragment ions and their correlations to assign glycan motifs^11^. Because this approach is analytically distinct from the precursor composition assignment approach performed by GlyComboCLI, the published results provide an orthogonal reference for validation. We therefore compared whether GlyComboCLI recovered the glycan motifs and tissue-specific trends reported in the original study.

The local pipeline, covering mzML conversion, GlyComboCLI, and Skyline, was applied to 52 raw files to identify and quantify glycan compositions (**Fig 2A**). Parallelisation ensured near-complete system resource utilisation, and all steps except verification were completed within 3 hours (**Fig 2B**). Beyond the automatic steps, which accounted for 60% of the total workflow duration, a 60-minute refinement step within Skyline was performed to ensure quality control and obtain quantitative values. Specifically, quality control was performed to reduce false identifications while retaining true positives for subsequent verification steps.

**Figure 2.**
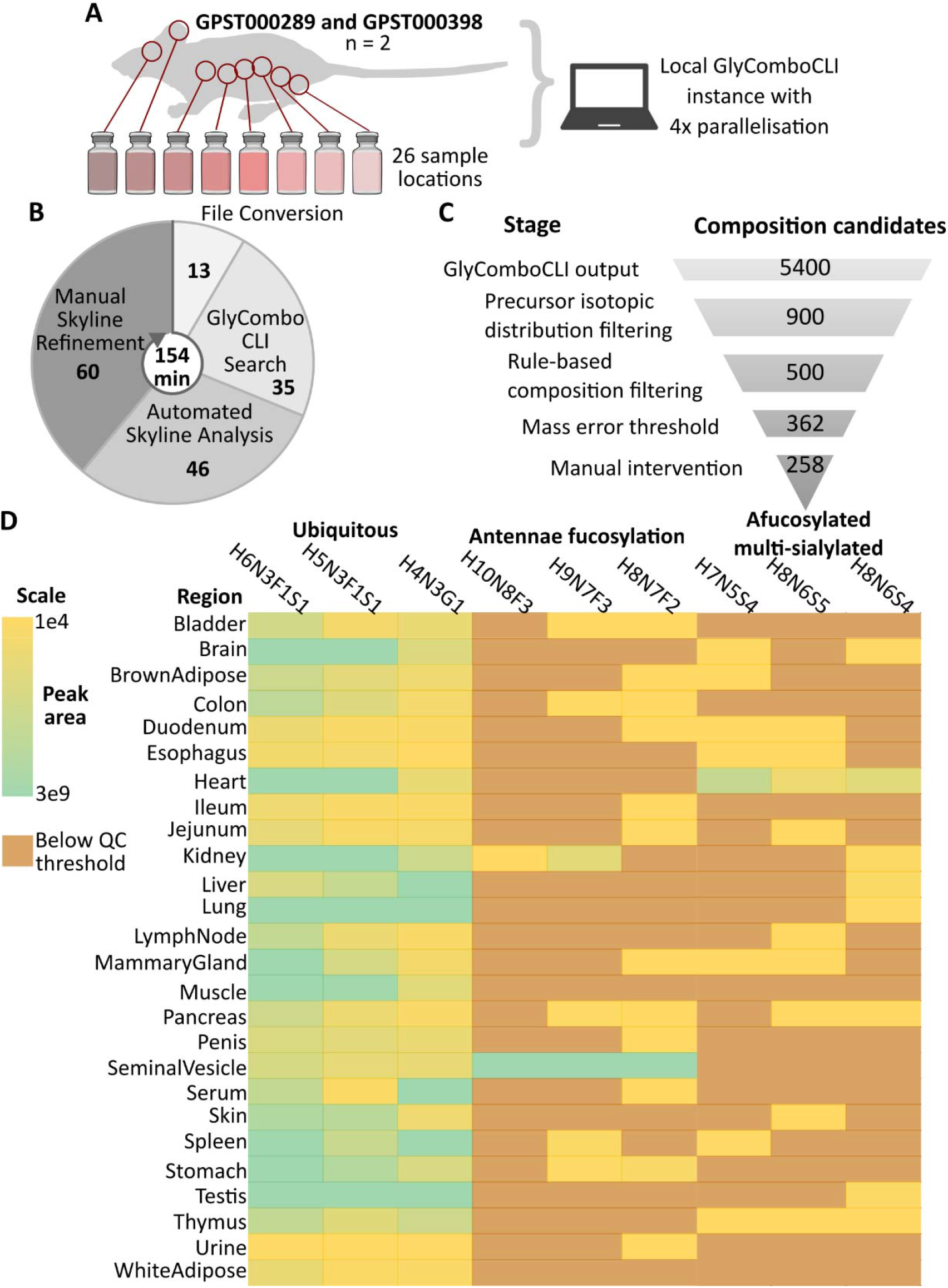
A local GlyComboCLI-enabled pipeline demonstrates high-throughput analysis of and orthogonal validation using a large-scale glycomics dataset. **A** Data summary and bioinformatic workflow used. **B** Time elapsed for each step in the bioinformatic workflow. **C** Filtering steps involved in generating high quality glycan composition assignments and effect on candidate numbers. **D** Comparison of GlyComboCLI-derived glycan features and tissue-associated trends with those reported in the original study.

As GlyCombo assigns compositions based on the combinatorial assignment of monosaccharides to precursor *m/z* values subjected to MS2, precursor *m/z* measurement is inherent and refinement is necessary. Following GlyComboCLI assignment, downstream QC reduced the candidate set substantially (**Fig 2C**). Skyline isotopic-envelope assessment removed poor-quality precursor matches and monoisotopic assignment errors, script-based composition filtering removed unlikely candidates (NeuAc/NeuGc ≤ Hex – 3, Supplementary Code), and a final 15 ppm mass error threshold was applied for quantitation.

Comparison with the original study showed concordance across several glycan motifs (**Fig 2D**). GlyComboCLI similarly identified hybrid *N*-glycan compositions (H6N3S1, H1H5N3S1, and H4N3G1) across all samples, antennae fucosylation, originally highlighted as a tissue-specific motif for seminal vesicles, was similarly identified in our approach (H10N8F3, H9N7F3, and H8N7F2), and a notable subset of glycan compositions, simultaneously absent of fucose (afucosylated) and multiply sialylated, were greatly enriched in the heart tissue regions. Agreement was therefore assessed at the level of reproducible glycan features and tissue-associated trends rather than as a one-to-one comparison of assignments, reflecting the distinct composition- and fragment-based analytical strategies used by the two approaches.

To complement the validation derived from the large and heterogeneous mouse tissue dataset, GlyComboCLI was additionally evaluated against a previously published human serum *N*-glycan dataset^35^ (**Figure 3**). GlyComboCLI assigned 168 putative glycan compositions, which were reduced to 105 following isotopic-envelope scoring and filtering of composition assignments. Comparison to the originally published identifications demonstrated substantial qualitative agreement, with a 69% overlap in composition identifications (**Fig 3A**). Quantitative reproducibility was further assessed by comparing reported abundance ranks, derived from the maximum glycan intensities reported in the original study, with those obtained using GlyComboCLI MS2 scan total ion current (TIC), and GlyComboCLI followed by Skyline MS1 peak area integration (**Fig 3B**).

**Figure 3.**
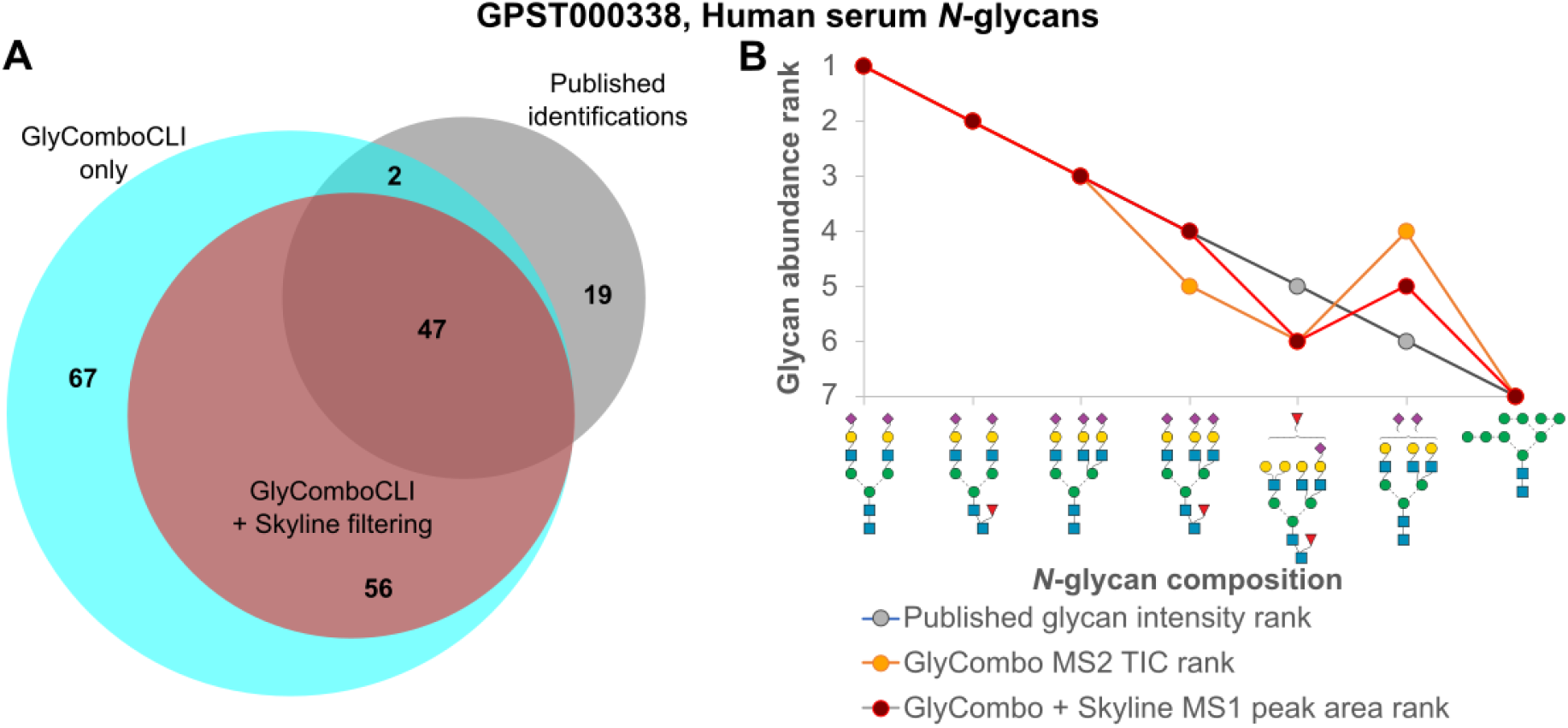
GlyComboCLI reproduces qualitative and quantitative features of a published human serum *N*-glycome. **A** Venn diagram comparing glycan compositions reported in the original study with GlyComboCLI assignments following isotopic-envelope filtering. **B** Rank trajectory plot comparing glycan abundance ranks derived from published glycan intensities, semi-quantitative GlyComboCLI MS2 scan total ion current, and Skyline MS1 peak-area integration.

MS2 TIC-based ranking was included as an easily automated, single scan-level comparison of the product ion signal, rather than as an ideal measure of glycan abundance. MS2 TIC is influenced by precursor isolation, fragmentation efficiency, and other acquisition parameters representing discrete sampling events while MS1 peak areas represent continuous precursor sampling across chromatographic elution, offering more appropriate measurements for relative precursor abundance comparisons. Despite these differences, the three most abundant glycans were ranked identically by all approaches. Some rank variation was observed for glycans ranked fourth to sixth, whereas H9N2 was consistently the lowest ranked glycan. The agreement between published abundance rankings and Skyline MS1 peak areas supports the reproduction of major relative abundance trends by GlyComboCLI while the MS2 TIC comparison illustrates the potential utility of MS2-level signal ranking.

### A reproducible approach to glycan composition assignment

Utilising publicly available online datasets ensures analytical consistency and provides accessible benchmarks for future software evaluation. Leveraging glycomics data from repositories such as GlycoPOST^36^ and Panorama Public^27^, files can be directly downloaded to a web server and processed remotely.

One such remote service is Galaxy, an open-source web-based platform designed to improve research accessibility, reproducibility and transparency. Galaxy operates through connected data flows between individual tools and accordingly, GlyComboCLI was prepared for the Galaxy ToolShed by wrapping a Debian-based Docker image of GlyComboCLI in a Galaxy XML wrapper. In addition to specifying inputs, parameters, and outputs for the web interface, the Galaxy XML wrapper encodes 21 embedded functional tests covering text and mzML inputs, instrument-vendor-derived files, search parameters, and custom search functionality to ensure consistency across versions. Within Galaxy, msconvert, GlyComboCLI, and ggplot2 can be combined in a single workflow to generate standard GlyCombo output alongside QC plots and analysis summaries (**Fig 4**). This workflow is constrained by the tools available within Galaxy and therefore components of the local pipeline, such as Skyline, cannot be incorporated.

**Figure 4.**
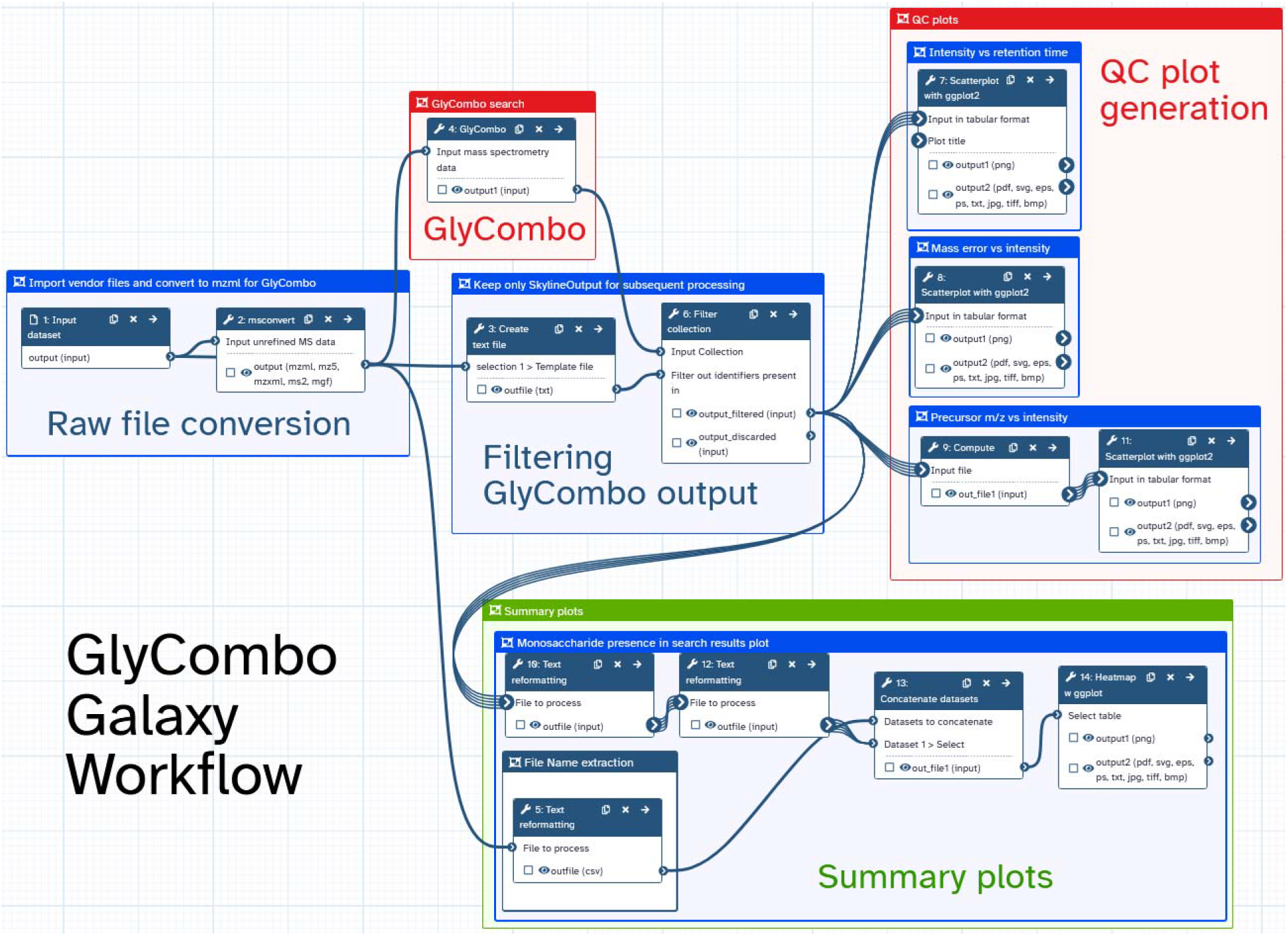
Visualisation of a glycomics Galaxy workflow that readily generates summary plots and QC plots from GlyComboCLI output

The Galaxy GlyCombo workflow was applied to sialidase treated data available on Panorama to evaluate the effects of sialic acid removal on reported glycan compositions (**Fig 5A**). Although MS2 counts remained reasonably constant, putative glycan identifications were reduced to 36% of those in the control sample (**Fig 5B**). A heatmap generated within the workflow summarised the GlyComboCLI output by comparing the intensity of the peaks assigned to compositions containing each monosaccharide. Compositions containing deoxyhexose (dHex), hexose (Hex), and *N*-acetylhexosamine (HexNAc) remained constant across both files, while NeuAc was reduced 10-fold, indicating effective sialic acid removal by sialidase (*Fig 5C*). Quality control plots were also generated to display the distribution of these glycan compositions across retention time and precursor *m/z* as compared to intensity, and mass error compared to intensity. The latter was used to verify that mass error tolerances for the GlyComboCLI search were appropriate (**Fig 5D**).

**Figure 5.**
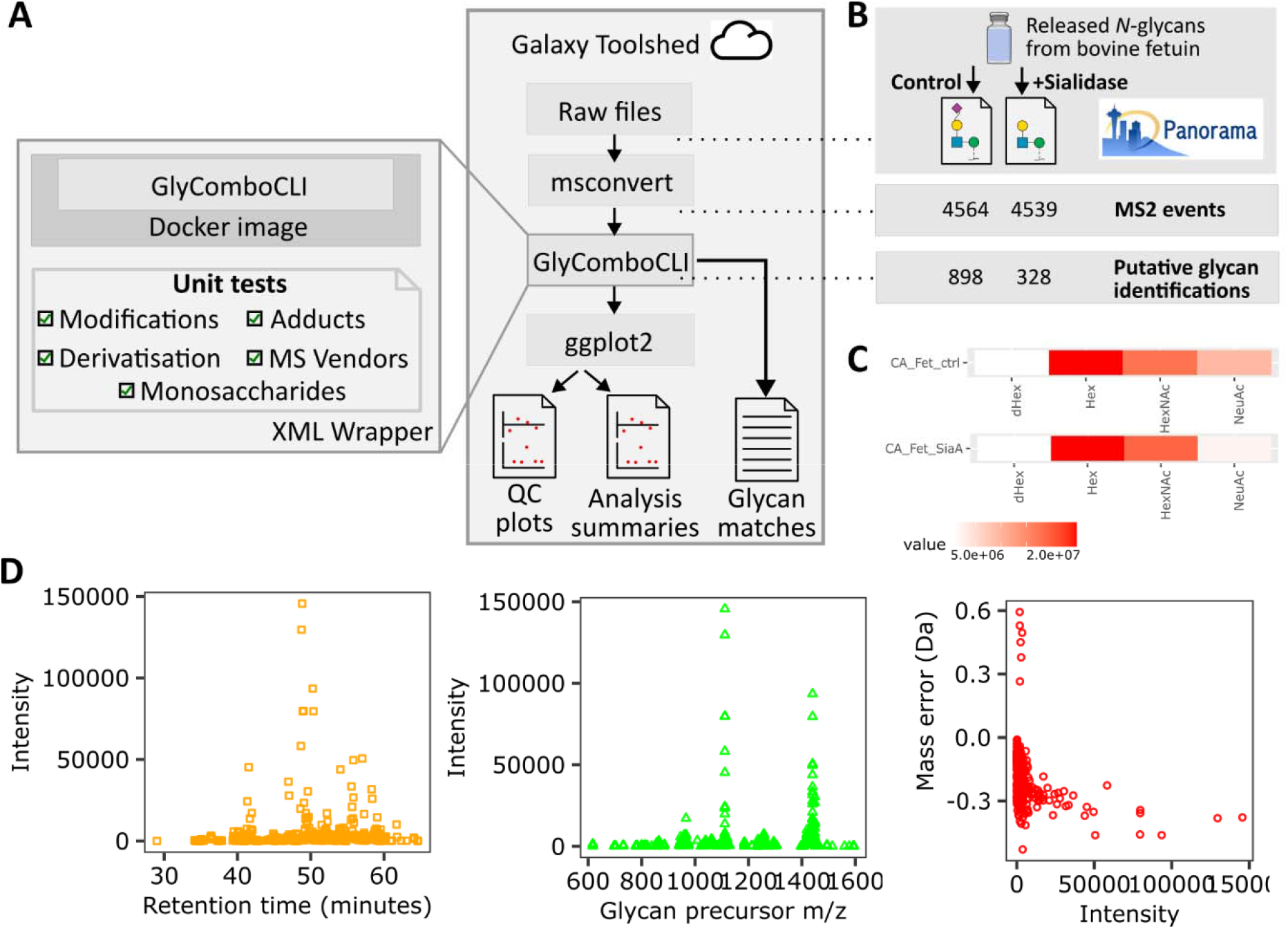
Glycan composition assignment workflow within Galaxy. **A** Galaxy retrieval of local or remote raw MS files for vendor-neutral conversion, glycan composition assignment, and visualisation. **B-D** Sialidase treatment subjected to the Galaxy glycomics workflow, revealing quantitative differences **(C)** and generating QC plots **(D)**.

## Discussion

The accessibility and efficiency of GlyComboCLI positions it well for widespread re-analysis of existing datasets such as those deposited on GlycoPOST. The 3-hour completion time across 52 files using parallelisation on a local computer is encouraging in that regard. As this is one of the largest GlycoPOST repositories, it bodes well for application across the >400 datasets made available in the entire repository. Validation across complementary published datasets demonstrates the reproducibility of GlyComboCLI across differing biological and analytical contexts. The mouse tissue analysis reproduced previously reported glycomic features, while the human serum benchmark showed substantial agreement in both composition identification and relative abundance trends. Although these comparisons do not constitute an absolute ground truth, they provide complementary evidence that GlyComboCLI generates results consistent with established analyses.

Direct software-to-software performance comparisons remain challenging because GlyComboCLI differs in analytical scope from existing tools, other than GlycoMod^13^. While related software such as GlycReSoft^37^, GlycoGenius^38^, and AnnotatoR^39^ address overlapping problems, they differ in scope as the former two incorporate peak picking and MS1-based annotation, while AnnotatoR focuses on MS1 annotation. GlycoGenius is designed to handle raw data, including peak picking and quantification, and it uses MS2 primarily to validate the initial MS1-based assignments. In contrast, GlyComboCLI is a command-line tool designed to be specifically concerned with assigning glycan compositions to precursor masses subjected to MS2. Benchmarking against these tools represents a clear opportunity for future development.

The extension and update of GlyCombo also ensures that the software supports FAIR principles. This is increasingly important as the global software ecosystem becomes more complex, and as automation becomes more important for life science workflows. GlyCombo, and its metadata, are easy for both machines and humans to **find** and understand. This is achieved using a combination of globally unique and persistent identifiers, including bio.tools software ID (https://bio.tools/glycombocli) , rich metadata that is both searchable and indexable, as well as DOIs for both the software itself, and the accompanying Galaxy workflow (https://doi.org/10.48546/WORKFLOWHUB.WORKFLOW.2167.1, https://doi.org/10.5281/zenodo.18736717).

The DOIs also make the software persistently **accessible**, and the bio.tools / WorkflowHub^40^ registry entries make the metadata available via a standardised protocol. GlyComboCLI also supports **interoperability**, meeting domain expectations through the use of vendor-neutral mzML input format and machine-readable outputs compatible with downstream glycoinformatics tools (*e*.*g*. GlycoWorkBench, GlyConnect, GlyGen), and GlyTouCan accessions for registered base compositions. These features facilitate exchange with the broader glycomics software ecosystem. **Reusability** is supported through open-source distribution under the Apache 2.0 license, documented local and containerised deployment, and incorporation into reproducible Galaxy workflows. Together, these features support alignment with FAIR principles.

Refinement of GlyCombo output is recommended given the potential for false positives, and full structural interpretation remains difficult to automate due to its dependence on analyte type, glycan derivatisation, and method of detection. These false positives typically arise from incorrect monoisotopic precursor assignment made in real-time by the mass spectrometer, whereas structural elucidation is performed using orthogonal evidence such as retention time, MS2 fragmentation spectra, and exoglycosidase or linkage-specific derivatisation treatments, which vary across the field and lack consensus^41^. As the Galaxy pipeline excludes Skyline, which we have used in our local pipeline to remove false positives and incorporate this orthogonal evidence, structural insight is limited. Regardless, GlyComboCLI substantially reduces the burden on expert users by narrowing the pool of MS2 spectra requiring structural consideration by up to 92% (**Figure 5B**), making subsequent manual interpretation more tractable.

Further improvements to performance and accessibility are possible. For example, incorporating sample preparation metadata to guide parameter selection, and automatically including monosaccharide compositions used by N-glycans, if PNGase-F treatment is recorded, to reduce the configuration burden on users and improve search relevance. Additionally, a survey search could be implemented to inform and refine search parameters prior to a full run, such as adjusting mass error tolerances and excluding monosaccharides not detected in the preview, improving the throughput and specificity of the final output.

## Conclusions

GlyComboCLI addresses a practical gap in glycoinformatics by enabling glycan composition assignment within automated, reproducible workflows across a range of deployment environments. Benchmarking against published mouse tissue and human serum N-glycan datasets further demonstrated that GlyComboCLI can reproduce major qualitative glycomic features and relative abundance patterns across analytically distinct datasets.

Its compatibility with existing community tools and adherence to FAIR principles facilitates integration into both local and shared computational pipelines, lowering barriers for researchers with varying levels of bioinformatic expertise. While structural interpretation remains a largely manual process dependent on orthogonal evidence, GlyComboCLI meaningfully reduces the volume of spectra requiring expert attention. Future developments, such as incorporation of sample preparation metadata and adaptive search parameters, could further improve usability and search relevance. The availability of GlyComboCLI as an open-source tool with documented workflows also supports re-analysis of publicly deposited glycomics datasets, contributing to the broader goal of reproducible glycomics research.

## Data Availability

The open-source GlyComboCLI code, under the Apache 2.0 license, is publicly available on GitHub at https://github.com/Protea-Glycosciences/GlyComboCLI and archived on Zenodo (https://doi.org/10.5281/zenodo.20063128), with software metadata registered at bio.tools (https://bio.tools/glycombocli). Versioned GlyComboCLI container images are available from Docker Hub at https://hub.docker.com/r/proteaglycosciences/glycombo. The Galaxy wrapper source is available at https://github.com/Protea-Glycosciences/GlyComboGalaxy and the validated Galaxy tool is distributed through the Main Galaxy Tool Shed at https://toolshed.g2.bx.psu.edu/repos/proteaglycosciences/glycombo. The Galaxy workflow is archived at https://doi.org/10.48546/WORKFLOWHUB.WORKFLOW.2167.1.

## Supplementary Data statement

GlyComboCLI output schematic (**Figure S1; PDF**); annotated MS2 spectra supporting compositions described in Figure 2 (PDF); composition assignments and quantitative data underlying the reproducibility assessment in Figure 3 (XLSX).

## SUPPORTING INFORMATION

Figure S1. GlyComboCLI output structure and representative result examples

Figure S2. Annotated MS2 spectrum of (Hex)6 (HexNAc)3 (NeuAc)1 from mouseNglycome_brain_2.raw at scan # 7315

Figure S3. Annotated MS2 spectrum of (Hex)5 (HexNAc)3 (dHex)1 (NeuAc)1 from mouseNglycome_brain_2.raw at scan # 7931

Figure S4. Annotated MS2 spectrum of (Hex)9 (HexNAc)6 (dHex)2 from mouseNglycome_seminalVesicle_2.raw at scan # 11129

Figure S5. Annotated MS2 spectrum of (Hex)3 (HexNAc)3 (dHex)3 from mouseNglycome_seminalVesicle_1.raw at scan # 7096

Figure S6. Annotated MS2 spectrum of (Hex)9 (HexNAc)7 (dHex)3 from mouseNglycome_seminalVesicle_2.raw at scan # 13352

Figure S7. Annotated MS2 spectrum of (Hex)8 (HexNAc)6 (NeuAc)5 from mouseNglycome_heart_1.raw at scan # 9999

Figure S8. Annotated MS2 spectrum of (Hex)6 (HexNAc)3 (dHex)1 (NeuAc)1 from mouseNglycome_brain_2.raw at scan # 8335

Figure S9. Annotated MS2 spectrum of (Hex)8 (HexNAc)6 (NeuAc)4 from mouseNglycome_heart_1.raw at scan # 10270

Figure S10. Annotated MS2 spectrum of (Hex)7 (HexNAc)5 (NeuAc)4 from mouseNglycome_heart_1.raw at scan # 9787

Data S1. SupplementaryData.xlsx contains the Figure 3 supporting information for publication.

Data S2. Supplementary_Code.zip contains the script-based composition filtering to remove unlikely candidates

## Funding/acknowledgments

The authors acknowledge Dr. Cameron Hyde for input and design into the Galaxy GlyComboCLI tool. Dr Hyde is employed by QCIF Pty Ltd. His contributions to Galaxy Australia are funded by NCRIS through Bioplatforms Australia, and The Queensland Government RICF fund.

O.J.R.G is supported by Australian BioCommons, which is enabled by NCRIS via Bioplatforms Australia funding. This work is supported by Galaxy Australia, a service provided by Australian BioCommons and its partners. The service receives NCRIS funding through Bioplatforms Australia, as well as The University of Melbourne and Queensland Government RICF funding.

## For TOC Only

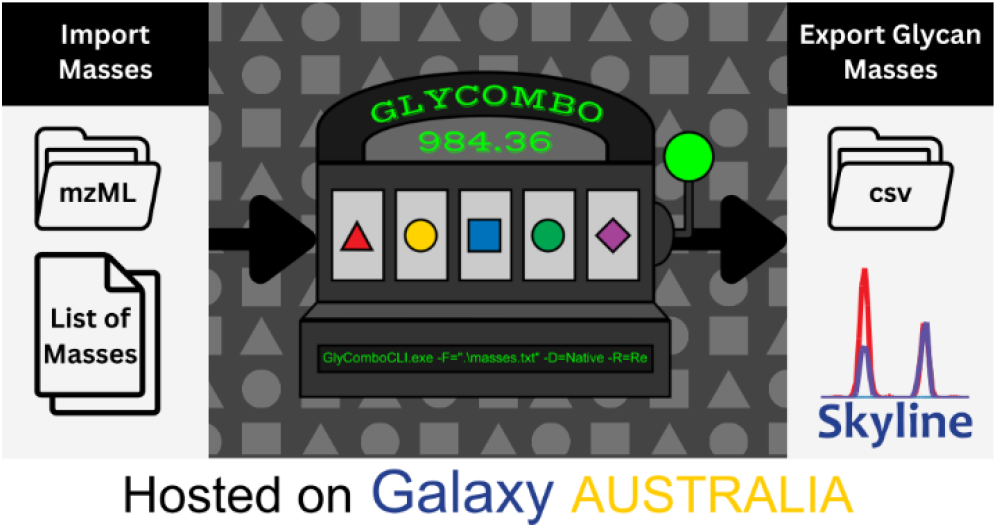

