## Supplementary material for "GlyComboCLI enables command line-based FAIR workflows for glycan composition assignment in mass spectrometry data": Figure S

### Supplementary figure and annotated spectra

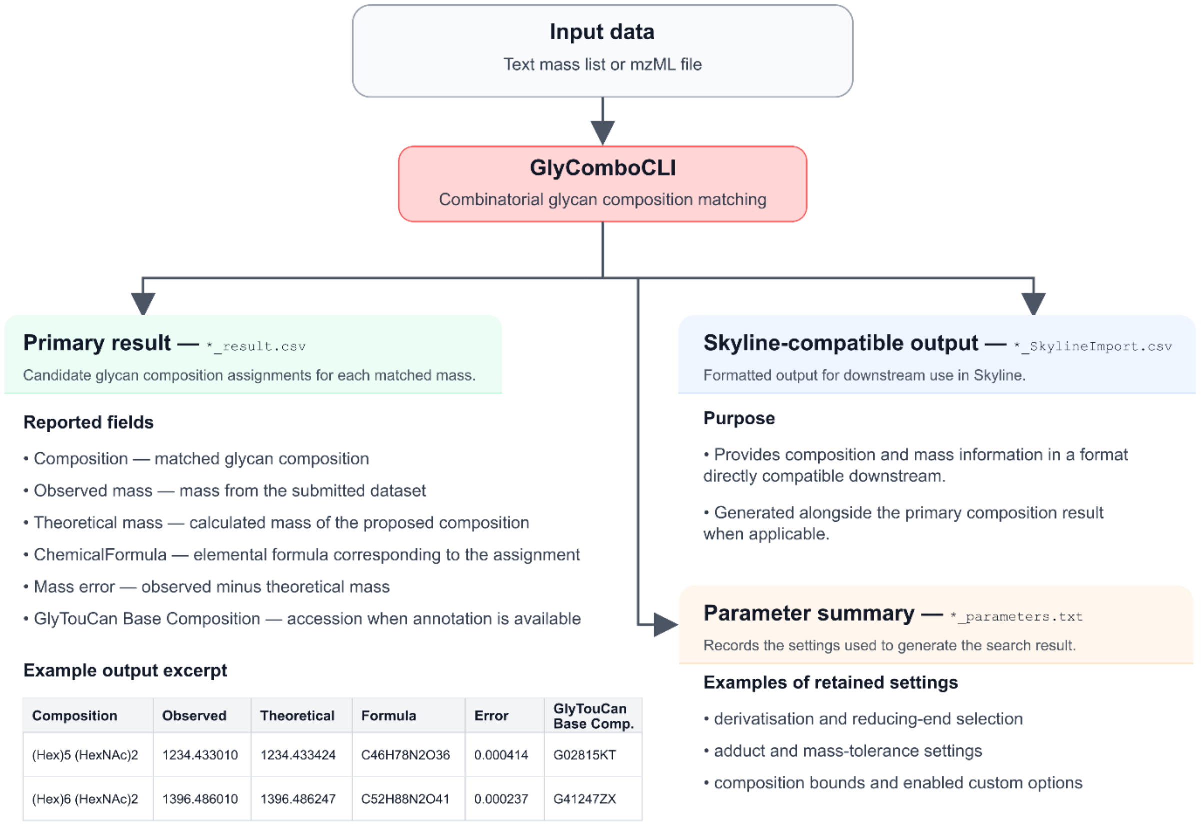

**Figure S1** GlyComboCLI output structure and representative result examples

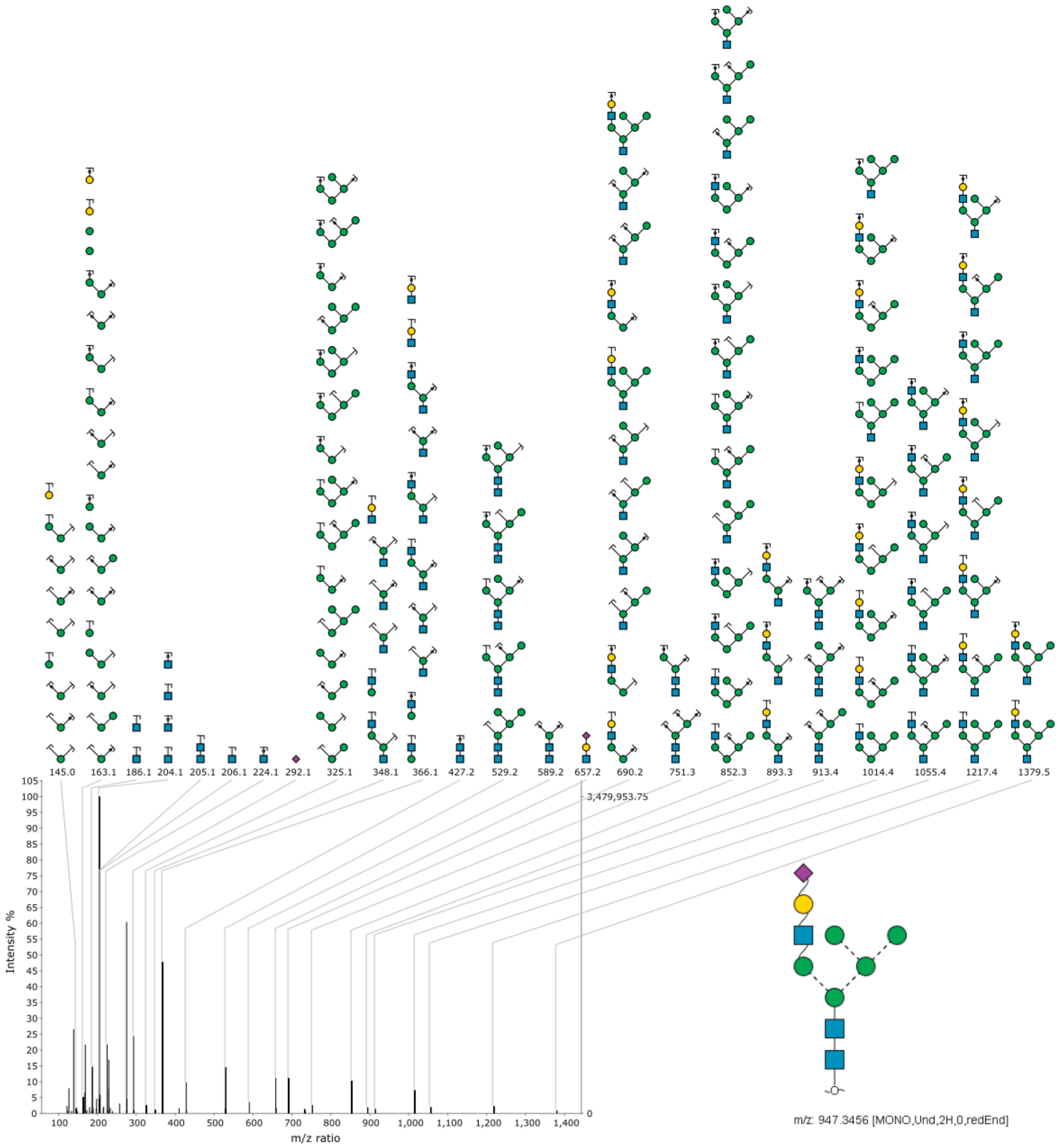

**Figure S2** Annotated MS2 spectrum of (Hex)6 (NeuAc)1 (HexNAc)3 from mouseNglycome_brain_2.raw at scan # 7315

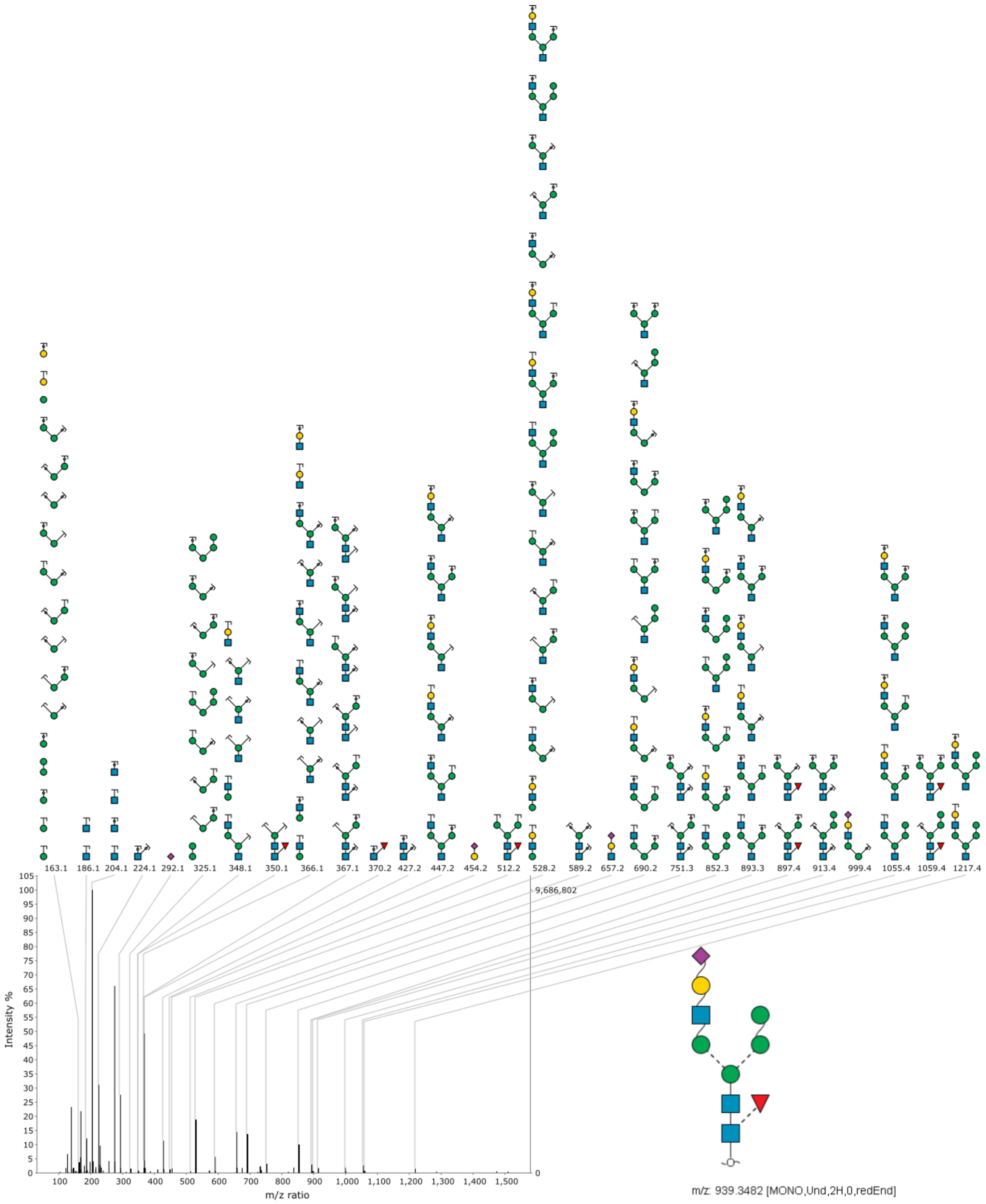

**Figure S3** Annotated MS2 spectrum of (dHex)1 (Hex)5 (NeuAc)1 (HexNAc)3 from mouseNglycome_brain_2.raw at scan # 7931

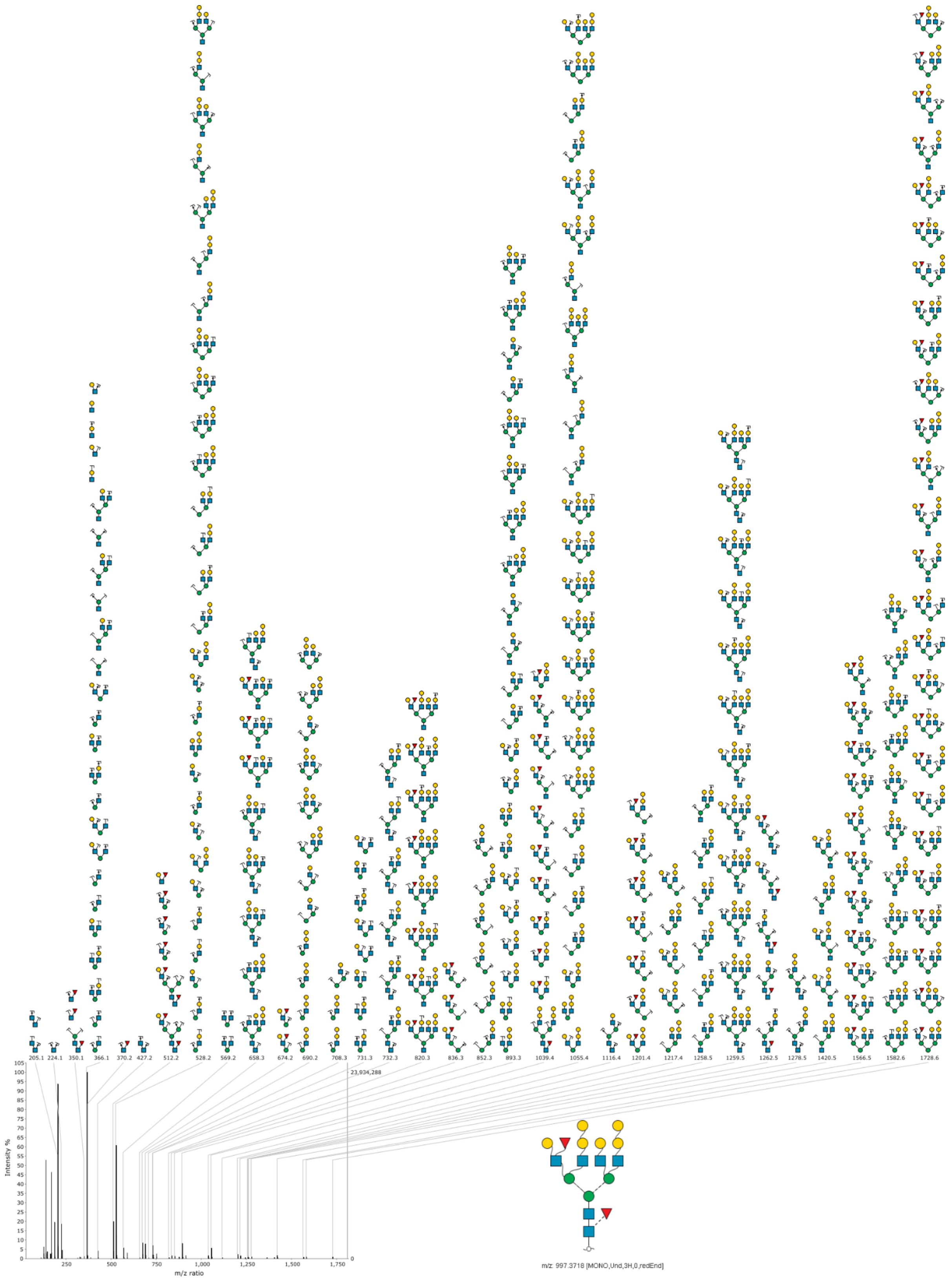

**Figure S4** Annotated MS2 spectrum of (dHex)2 (Hex)9 (HexNAc)6 from mouseNglycome_seminalVesicle_2.raw at scan # 11129

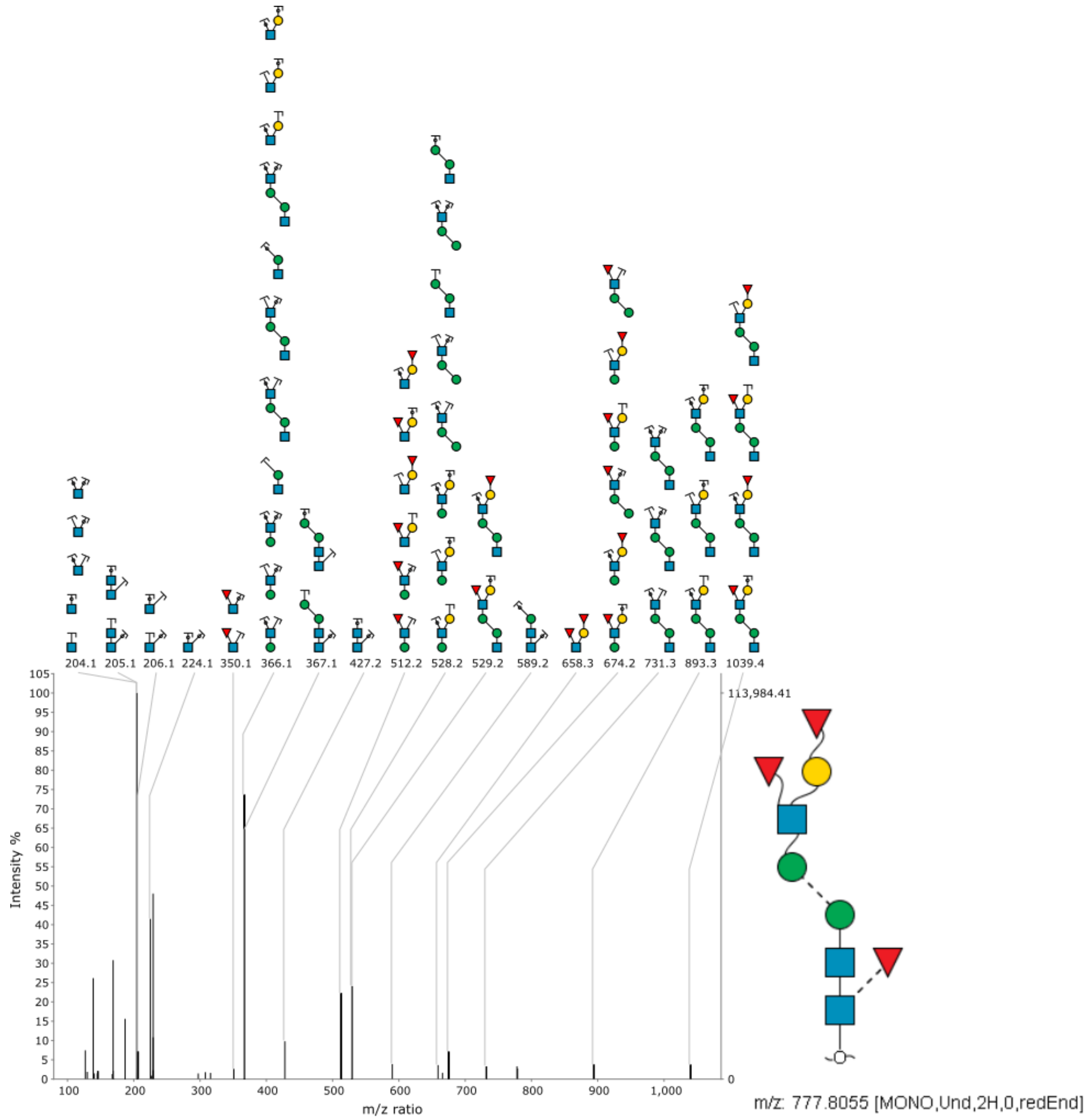

**Figure S5** Annotated MS2 spectrum of (dHex)3 (Hex)3 (HexNAc)3 from mouseNglycome_seminalVesicle_1.raw at scan # 7096

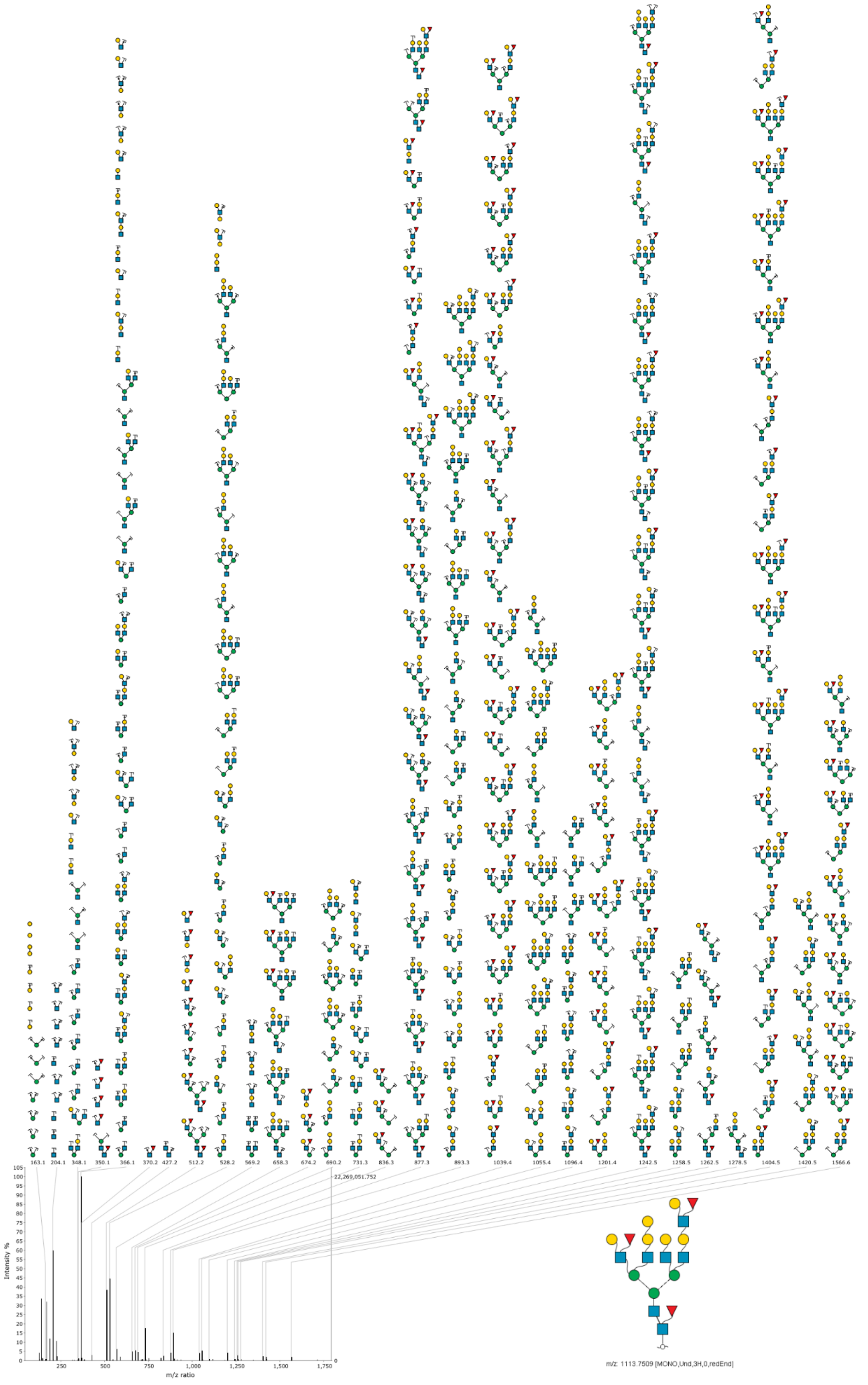

**Figure S6** Annotated MS2 spectrum of (dHex)3 (Hex)9 (HexNAc)7 from mouseNglycome_seminalVesicle_2.raw at scan # 13352

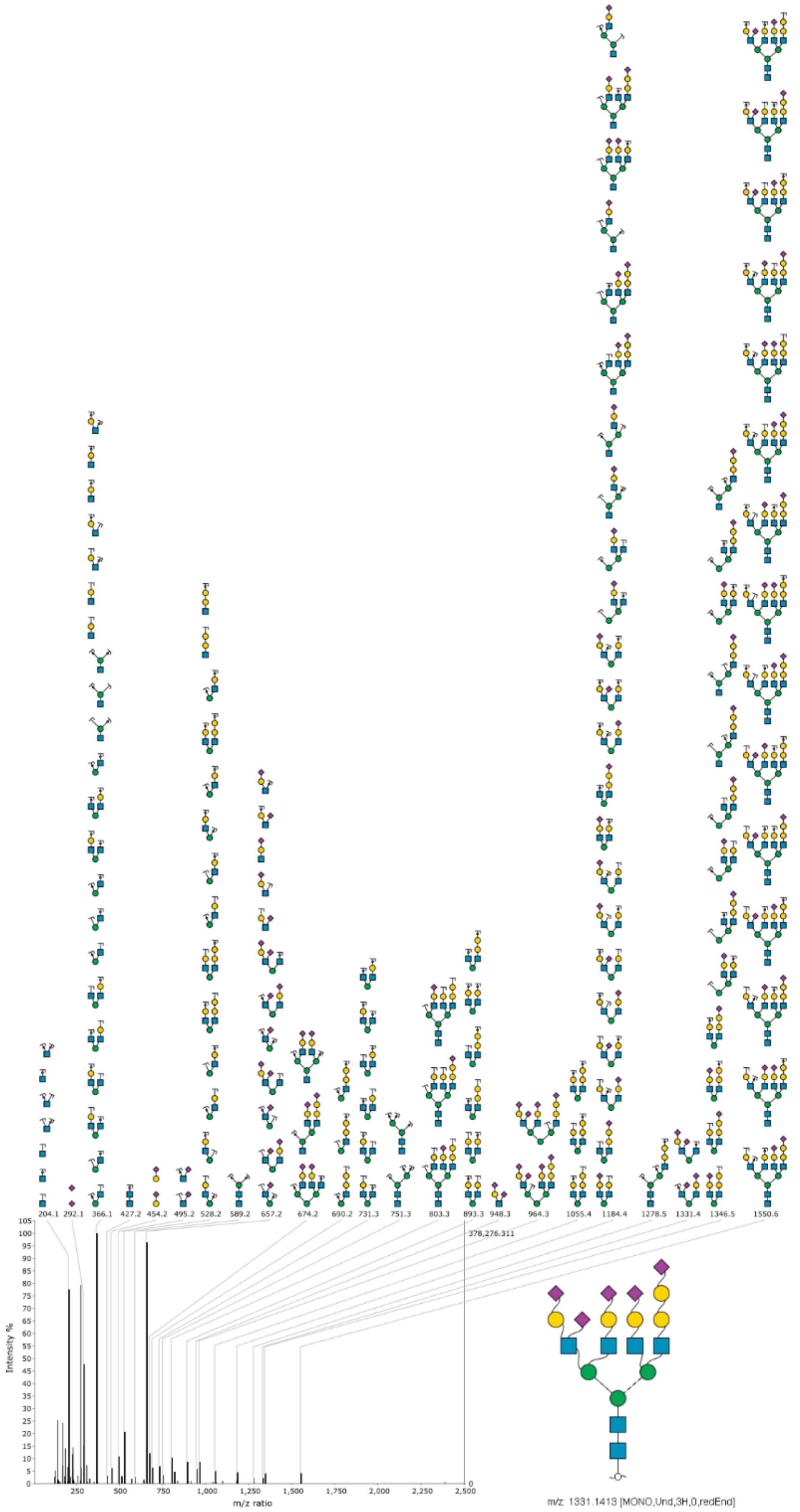

**Figure S7** Annotated MS2 spectrum of (Hex)8 (NeuAc)5 (HexNAc)6 from mouseNglycome_heart_1.raw at scan # 9999

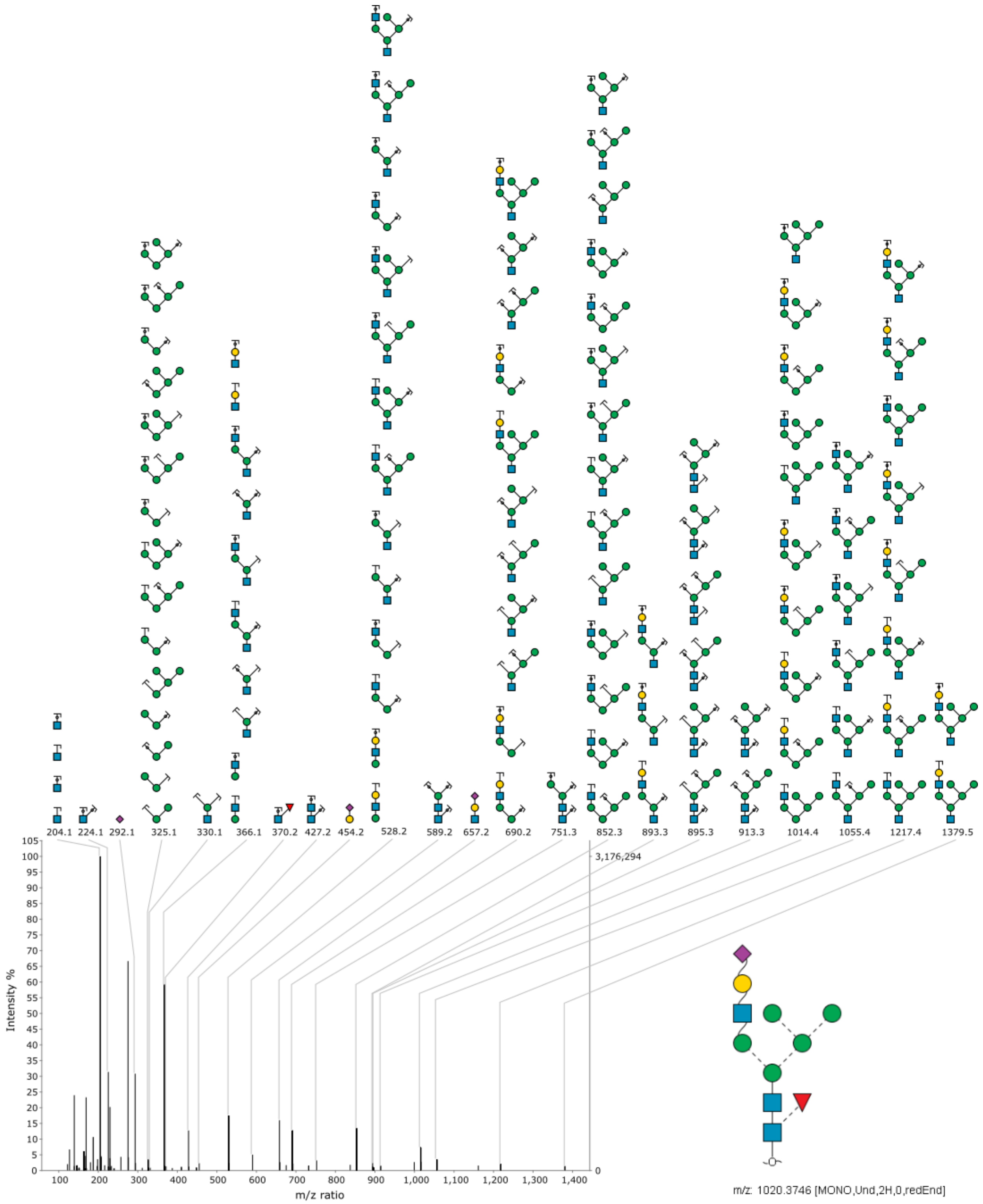

**Figure S8** Annotated MS2 spectrum of (dHex)1 (Hex)6 (NeuAc)1 (HexNAc)3 from mouseNglycome_brain_2.raw at scan # 8335

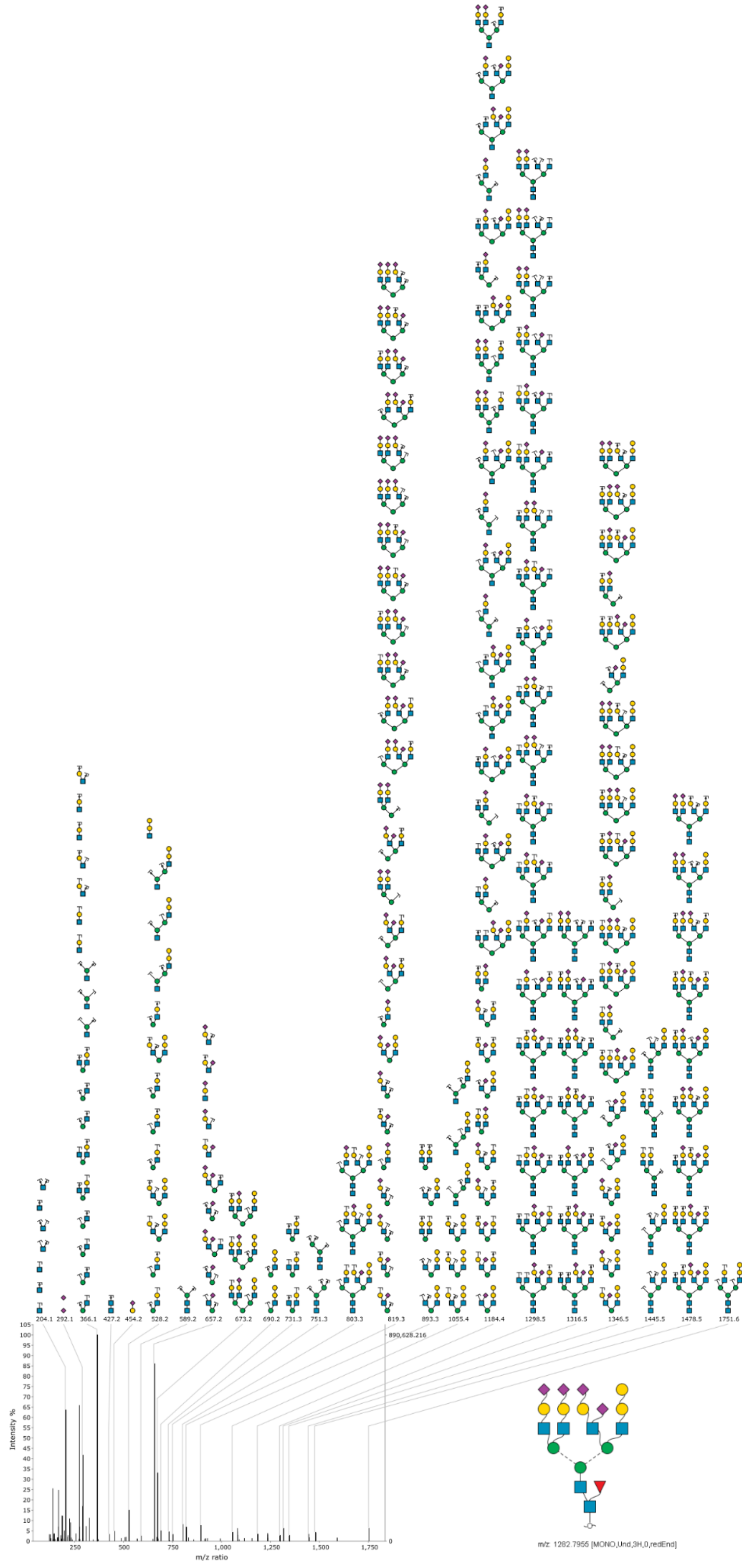

**Figure S9** Annotated MS2 spectrum of (Hex)8 (NeuAc)4 (HexNAc)6 from mouseNglycome_heart_1.raw at scan # 10270

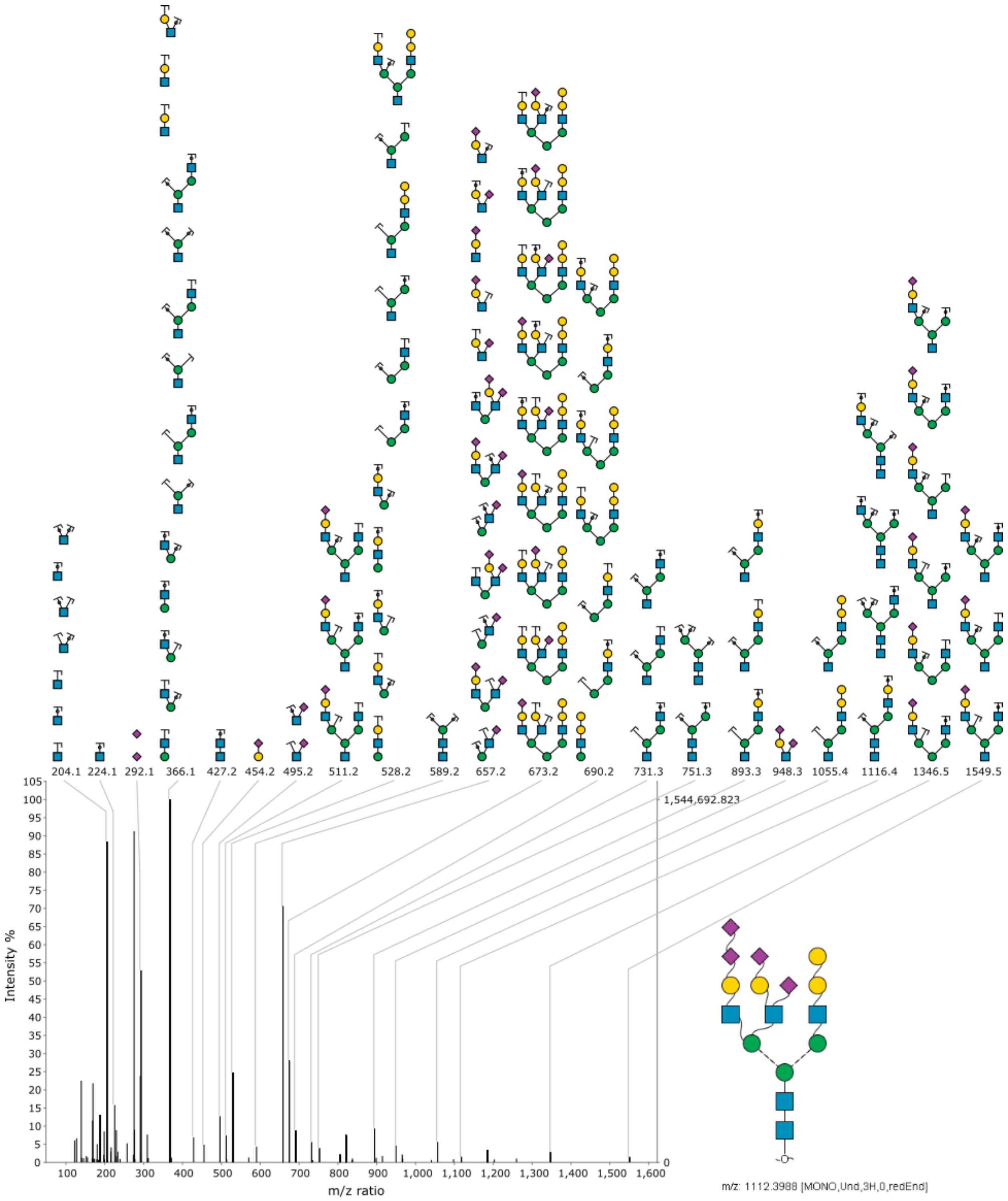

**Figure S10** Annotated MS2 spectrum of (Hex)7 (NeuAc)4 (HexNAc)5 from mouseNglycome_heart_1.raw at scan # 9787
